# Lower-limb rigidity is associated with frequent falls in Parkinson disease

**DOI:** 10.1101/515155

**Authors:** J. Lucas McKay, Madeleine. E. Hackney, Stewart A. Factor, Lena H. Ting

**Author notes:** These authors contributed equally to this work (J. Lucas McKay, Madeleine. E. Hackney). Corresponding author Mail: 1760 Haygood Drive NE, Room W-202, Atlanta, GA 30322, USA, (JLM).

## Abstract

**BACKGROUND AND OBJECTIVE:** The role of muscle rigidity as an etiological factor of falls in Parkinson disease (PD) is poorly understood. Our objective was to determine whether lower leg rigidity was differentially associated with frequent falls in PD compared to upper limb, neck, and total rigidity measures. METHODS: We examined associations between UPDRS-III (motor) rigidity subscores and history of monthly or more frequent falls in N=216 individuals with PD (age, 66±10 y; 36% female, disease duration, 7±5 y) with logistic regression. RESULTS: N=35 individuals were frequent fallers. Significant associations were identified between lower limb rigidity and frequent falls (P=0.01) after controlling for age, sex, PD duration, total UPDRS-III score, and presence of FOG. No significant associations (P≥0.14) were identified for total, arm, or neck rigidity. CONCLUSION: Lower limb rigidity is related to frequent falls in people with PD. Further investigation may be warranted into how parkinsonian rigidity could cause falls.

Financial Disclosures/Conflict of Interest concerning the research related to the manuscript: None

**Funding:** NIH K25HD086276, R01HD046922, R21HD075612, UL1TR002378, UL1TR000454; Department of Veterans Affairs R&D Service Career Development Awards E7108M and N0870W, Consolidated Anti-Aging Foundation, and the Sartain Lanier Family Foundation.

## Introduction

Balance problems and falls are a major issue in Parkinson’s disease (PD).^1^ Recently, a group of PD patients, caregivers, and health professionals ranked balance problems and falls as a primary research priority.^2^ While we have some ability to identify PD patients at high risk for falls based on factors like fall history,^3^ many factors are non-modifiable, and hence have little value in preventing falls. Although many studies have attempted to identify associations between clinical and physiological measures like including UPDRS^4^ items and falls,^3, 5, 6^ rigidity is an understudied factor.^5, 7, 8^

Parkinsonian rigidity may cause functional balance and gait impairments balance and gait,^9-12^ and responds well to levodopa,^9, 13-15^ but its relationship to falls is unclear. Biomechanical simulations^16^ suggest that rigidity, particularly lower limb rigidity, may be an important contributor to falls. In simulation, increased muscle stiffness like that identified in rigid patients^17^ impairs the ability to withstand perturbations to the center of body mass.^16^ The narrow stance common in PD may compensate for stiffness due to rigidity^16^ and related problems.^18^ Although some studies suggest a contribution of axial rigidity to balance impairments and fall risk,^19, 20^ lower limb rigidity is not commonly considered as an independent risk factor.^5-7^

Here, we used logistic regression to determine whether lower limb rigidity assessed on the UPDRS-III^4^ was associated with a history of monthly or more frequent falls in individuals with mild-moderate PD. We reasoned that lower limb rigidity would be more associated with fall history than upper limb or neck rigidity due to the lower limbs’ involvement in locomotion and static balance. We hypothesized that: 1. Lower limb rigidity scores would be associated with falls; 2. Upper limb, neck, and total rigidity scores would not be associated with falls.

## Methods

### Data sources

We used existing measures of N=216 PD patients from observational and rehabilitative studies we conducted from 2011–2015. Participants provided written informed consent according to protocols approved by the Institutional Review Boards of Emory University and the Georgia Institute of Technology. All participants met diagnostic criteria for PD according to either modified UK Brain Bank criteria^21^ or diagnostic criteria for “clinically definite” PD.^22^ Exclusion criteria included advanced stage dementia and inability to walk ≥3 meters with or without assistance. Beginning with N=220 records, 4 records were excluded due to incomplete Freezing of Gait questionnaire (FOG-Q)^23^ or UPDRS-III^4^ scores.

### Study variables

Participants were classified as “Fallers” if they scored ≥2 on Gait and Falls questionnaire (GF-Q) item 12,^23^ corresponding to monthly or more frequent falls, and were classified as “Non-Fallers” otherwise. The primary independent variables were rigidity subscores assembled from rigidity items of the UPDRS-III:^4^ total rigidity (/20), the sum of UPDRS-III items 22a-e; lower limb sum (/8), the sum of items 22d-e; upper limb sum (/8), the sum of items 22b-c; and neck (/4), item 22a. Additional demographic and clinical variables associated with falls in PD included age, female sex, global cognition (assessed with the Montreal Cognitive Assessment, MoCA^24^), disease duration,^3, 25^ and presence of freezing of gait (FOG). Participants were classified as freezers if they scored ≥2 on FOG-Q^23^ item 3, corresponding to weekly or more frequent freezing.

### Statistical methodology

Differences in study variables between Fallers and Non-Fallers were assessed with *t*-tests and chi-squared tests as appropriate. Satterthwaite’s approximation was used for *t*-tests with unequal variance as assessed with Folded F tests. Multivariate logistic regressions were performed to identify associations between rigidity subscores and faller status while controlling for age, gender, PD duration, total UPDRS-III score, and presence of FOG.^25^ Associations between rigidity subscores and fall history were expressed as Odds Ratios (OR) +/-95% Confidence Intervals. To control for overall UPDRS-III score, the remainder of each UPDRS-III total score after subtracting rigidity items was entered as a covariate. Statistical tests were performed at alpha ≤ 0.05 in SAS University Edition 7.2.

### Additional analyses

Additional analyses examined: associations between rigidity subscores for each limb and fall history, associations between rigidity subscores and annually or more frequent (rather than monthly or more frequent) falls, the impact of postural instability (UPDRS-III item 30), gait impairment (UPDRS-III item 29), overall cognition (MoCA score), and medication state during examination on associations between rigidity subscores and fall history (S1 File).

## Results

Demographic and clinical characteristics are presented in Table 1. Among the study sample, 35/216 (16%) fell monthly or more often. Consistent with previous results,^3, 25^ compared to Non-Fallers, Fallers had longer disease duration (P<0.01) and worse performance on UPDRS-III (P<<0.01), GF-Q (P<<0.01) and FOG-Q (P<<0.01). Among Fallers, prevalence of FOG and female sex were also increased by ≈3 times (P<<0.01) and ≈1.5 times (P=0.04), respectively. No significant differences were observed in age, global cognition (MoCA), education, or age at PD onset.

**Table 1.** Demographic and clinical features of the study population overall and stratified on presence of previous monthly falls.

| Characteristic | All Participants<br>N=216 | Non-Fallers<br>N=181 | Fallers<br>N=35 | P value |
| --- | --- | --- | --- | --- |
| Age, y | 65.7±9.7 | 65.5±9.6 | 67.1±10.3 | 0.35 |
| Sex |  |  |  |  |
| Female | 78 (36) | 60 (33) | 18 (52) | 0.04 |
| Male | 138 (64) | 121 (67) | 17 (48) |  |
| MoCA (/30) | 24.7 ± 3.6<sup>a</sup> | 24.8 ± 3.6<sup>b</sup> | 24.2 ± 4.0<sup>c</sup> | 0.37 |
| Education, y | 16.1 ± 2.2<sup>d</sup> | 16.1 ± 2.3<sup>e</sup> | 16.2 ± 1.7 | 0.92 |
| Disease duration, y | 7.4 ± 4.5 | 6.9 ± 4.1 | 9.9 ± 5.7 | <0.01 |
| Age at onset, y | 58.3 ± 10.6 | 58.6 ± 10.1 | 57.3 ± 12.9 | 0.58 |
| UPDRS-III score (/108) | 22.0 ± 10.0 | 20.6 ± 9.2 | 29.5 ± 10.5 | <<0.01 |
| FOG-Q Total | 4.5 ± 4.6 | 3.5 ± 3.9 | 9.6 ± 4.9 | <<0.01 |
| FOG-GF Total | 8.6 ± 9.0 | 6.1 ± 6.2 | 21.1 ± 10.9 | <<0.01 |
| Freezing of Gait |  |  |  |  |
| Freezer | 59 (27) | 35 (19) | 24 (69) | <<0.01 |
| Nonfreezer | 157 (73) | 146 (81) | 11 (31) |  |

Abbreviations: PD, Parkinson's disease; MoCA, Montreal Cognitive Assessment; UPDRS-III, Unified Parkinson's Disease Rating Scale, Part III: Motor Exam; FOG-Q, Freezing of Gait Questionnaire; FOG-GF, Gait and Falls Questionnaire. <sup>a</sup>N=194. <sup>b</sup>N=163. <sup>c</sup>N=31. <sup>d</sup>N=215. <sup>e</sup>N=180. Values are shown as either mean ± SD or N (%). P values reflect univariate tests of central tendency (*t*-tests or chi-squared tests) between Fallers and Non-Fallers.

Univariate analyses demonstrated that rigidity subscores for the lower limbs were significantly higher in Fallers (P value range: 0.004–0.025). In contrast, no statistically-significant differences were found for upper limb (P value range: 0.193–0.245) or neck (P=0.085) rigidity scores (Fig 1A).

**Figure 1.**
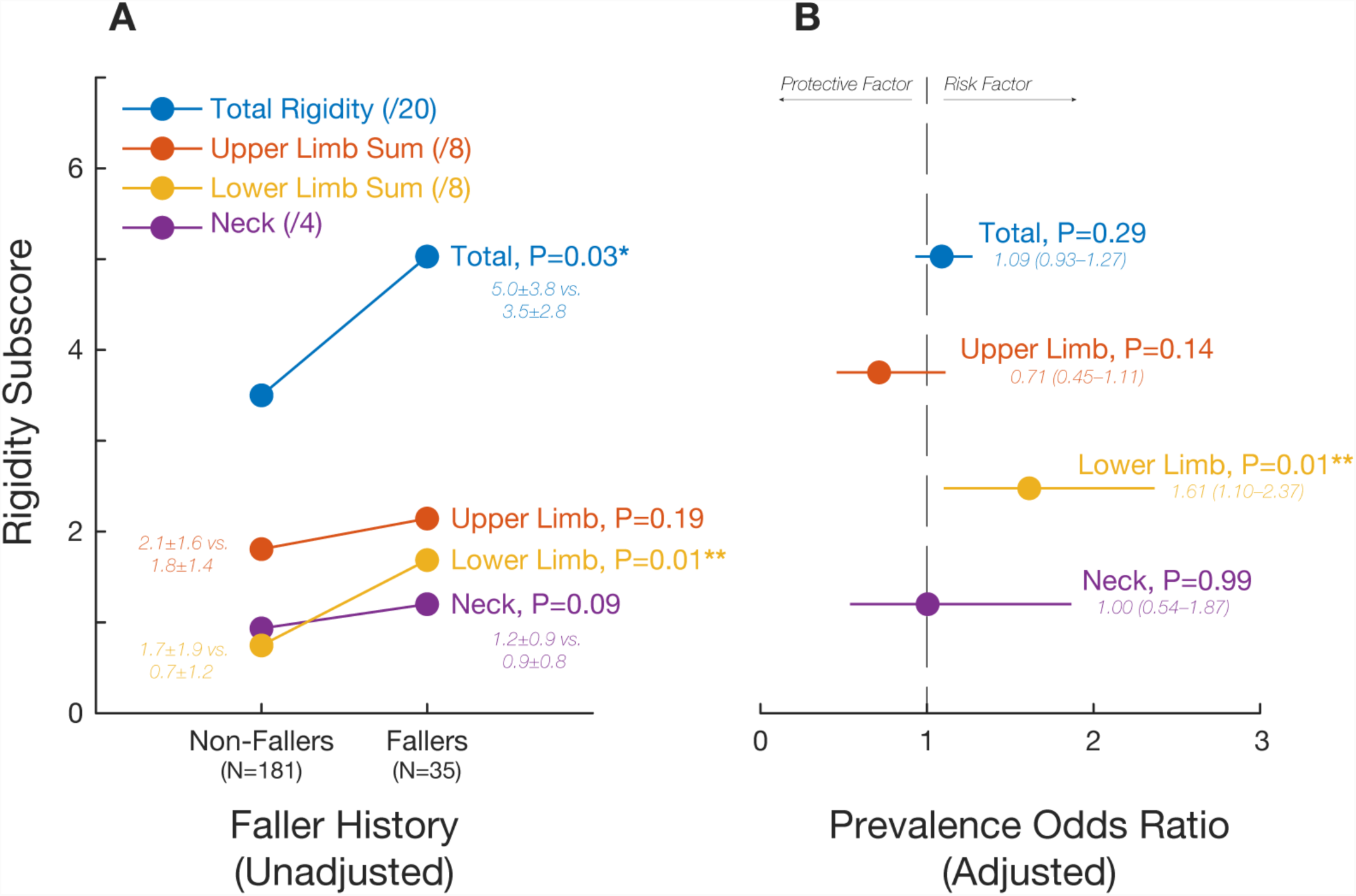
Associations between rigidity and falls. A) Differences in rigidity subscores between PD patients with and without history of monthly or more frequent falls. B) Associations between rigidity subscores and fall history (expressed as odds ratios [OR]±95%CI) after adjusting for age, sex, total UPDRS-III, PD duration, and presence of FOG.

Multivariate logistic regression models identified significant associations between lower limb rigidity and fall history (P=0.014; Fig 1B), but no significant associations for total (P=0.289), arm (P=0.135), or neck rigidity (P=0.991) after controlling for age, gender, PD duration, total UPDRS-III score, and presence of FOG. In these models, a total lower limb rigidity score of 2 (the 75^th^ percentile score in the sample) was associated with an odds ratio for monthly or more frequent falls of 2.6 (95% CI: 1.2–5.6), and a total score of 6 (the maximum score in the study sample), was associated with an odds ratio of 17.7 (1.8–175.7). Consistent with the literature,^3, 25^ logistic regression models also identified significant associations between female sex (Odds Ratio [95% CI] 3.2 [1.3–8.1], P=0.013), FOG (7.2 [2.9–17.9]; P<<0.01), disease duration (1.1[1.0–1.2]; P=0.022), and total UPDRS score (1.1 [1.0–1.2]; P=0.003) and fall history.

### Additional analyses

Associations between lower limb rigidity and history of frequent falls were largely unaltered in additional analyses (S1 File) that stratified participants by medication state during examination, or that controlled for the presence of postural instability (UPDRS item 30^5^), gait impairment (UPDRS item 29), or reduced cognition (MoCA score). These models yielded OR very similar to those identified in the main model, with changes in lower limb rigidity OR of -4.3%, +3.7%, - 8.1%, and +4.3%, respectively. When a less stringent definition of fall history (annual or more frequent rather than monthly or more frequent falls) was used, associations between lower limb rigidity and fall history were significantly reduced in magnitude (-21.7%), suggesting that lower limb rigidity is associated primarily with frequent falls. Models that included separate rigidity scores from each limb identified a strong association between rigidity on the more rigid side and history of frequent falls (OR increase of +23.6%).

## Discussion

To our knowledge, this is the first study to demonstrate an association between lower limb rigidity and falls in PD. We found that lower limb rigidity, unlike upper limb or neck rigidity, was associated with history of monthly or more frequent falls, even after controlling for common risk factors including age, sex, total UPDRS-III, PD duration, and presence of FOG. Additional analyses confirmed that the identified associations between lower limb rigidity and falls were not confounded by coexisting postural instability, gait impairment, or cognitive impairment in lower limb-rigid patients, as explicitly controlling for these factors altered odds ratios only minimally. These results suggest that understanding and treating rigidity may be important in reducing fall risk.

In addition to common features on exam that raise concerns to neurologists that falls may be impending for a patient, such as FOG, postural instability, and axial rigidity,^5, 6^ lower limb rigidity may be a clinically observable and potentially modifiable parkinsonian feature associated with falls. Notably, lower limb rigidity may be important to consider independent of axial rigidity,^9-11, 20, 26^ which is measured with specialized equipment^9, 11^ uncommon in clinical use. Although appendicular and axial rigidity are often thought of as having a common etiology, these signs likely reflect distinct underlying pathophysiology, and respond differently to dopaminergic drugs such as levodopa^9, 13, 15, 27^ and apomorphine.^14^

How might lower limb rigidity contribute to falls? Although abnormal deep tendon reflexes are not a key parkinsonian feature,^28^ rigid patients exhibit increased long latency electromyographic responses to passive joint movements,^29, 30^ and possibly increased tonic muscle activity,^31^ both of which may increase joint stiffness. A previous simulation study of standing balance^16^ demonstrated that as the stiffness of the hip joints are increased beyond a certain amount, the neuro-musculoskeletal system becomes increasingly unstable due to response delays. This makes it increasingly more difficult to maintain balance without stepping during perturbations. The implication is that a person with lower limb rigidity may have decreased ability to control the center of mass with the feet in place, requiring frequent steps to maintain balance. We hypothesize that because anticipatory postural adjustments^32, 33^ and stepping reaction time and accuracy are impaired in PD,^34^ lower limb rigidity could therefore cause falls. However, the underlying biomechanical and physiological mechanisms of rigidity remain unclear.

The primary limitation of this study is that fall history was taken via self-report. To limit this we used a stringent criterion to identify Fallers – monthly or more frequent falls. We found that the association between lower limb rigidity and falls was significantly attenuated in logistic regression models using a less stringent criterion. Based on this, we speculate that lower limb rigidity may be uniquely related with very frequent falls in PD; however, prospective studies are required to confirm this. If prospective studies demonstrate that lower limb rigidity is associated with future falls, patients with this sign could potentially be referred to interventions to reduce risk.^35-40^

Although many studies have attempted to identify associations between clinical and physiological assessments and fall risk;^3, 5, 6^ lower limb rigidity is an understudied risk factor.^5, 7^ This may be due in part to one impactful early study^6^ that found no association between the combined presence of lower limb / neck rigidity and fall risk, suggesting that combining these sources of rigidity may distort associations with falls. These results suggest that prospective studies of the relationships between rigidity and fall risk in PD could provide new information.

## Author roles

1) Research Project: A. Conception: JLM, LHT, MEH, SAF; Organization: MEH, JLM, LHT, SAF; C. Execution: MEH, JLM, LHT, SAF. 2) Statistical Analysis: A. Design, JLM B. Execution, JLM C. Review and Critique: MEH, SAF, LHT. 3) Manuscript: A. Writing of the first draft, MEH, LHT, JLM; B. Review and Critique: MEH, LHT, SAF.

## Supporting Information

S1 File. Supplementary Material. (PDF)

S2 File. Supporting Data. (CSV)

## Supplemental Material

### Associations between individual limb rigidity scores and monthly or more frequent falls

In addition to the primary rigidity subscores discussed in the main text, secondary analyses considered additional rigidity subscores calculated for individual limbs as follows: lower limb lesser (/4), the lesser of UPDRS-III^4^ items 22d-e; lower limb greater (/4), the greater of items 22d-e; upper limb lesser (/4), the lesser of items 22b-c; and, upper limb greater (/4), the greater of items 22b-c.

As in the main text, multivariate logistic regressions were performed to identify associations between secondary rigidity subscores and faller status while controlling for age, gender, PD duration, total UPDRS-III score, and presence of FOG.^25^ Associations between rigidity subscores and fall history were expressed as Odds Ratios (OR) +/-95% Confidence Intervals. In order to control for overall UPDRS-III score, the remainder of each UPDRS-III total score after subtracting the rigidity subscore of interest was entered as a covariate into logistic regression analyses. Prior to entry into multivariate analyses, age and disease duration were converted into z-scores.

Multivariate logistic regression results using individual limb rigidity subscores indicated that the association between lower limb rigidity and fall history described in the main text is primarily driven by a strong association between rigidity on the more affected side and falls. When individual limb rigidity scores were entered into models, we found that rigidity in each of the lower limbs – both the lesser and more affected sides – were positively associated with fall history, although the identified associations were no longer statistically significant (Table S2). In particular, the magnitude of association for the more affected side was increased (24%), whereas the magnitude of association for the less affected side was reduced (23%). This demonstrates that rigidity in the less-affected limb is much less strongly associated with fall history. Changes in associations among the upper limbs were minimal (2.8%).

**Table S1.** Rigidity subscores assembled from sub-items of the UPDRS-III motor scale, overall and stratified on presence of previous monthly falls.

| Rigidity Subscore | All Participants<br>(N=216) | Non-Fallers<br>(N=181) | Fallers<br>(N=35) | P value |
| --- | --- | --- | --- | --- |
| Total rigidity (/20) | 3.7 ± 3.0 | 3.5 ± 2.8 | 5.0 ± 3.8 | 0.03 |
| Lower limb sum (/8) | 0.9 ± 1.4 | 0.7 ± 1.2 | 1.7 ± 1.9 | <0.01 |
| Lower limb lesser (/4) | 0.3 ± 0.6 | 0.3 ± 0.5 | 0.7 ± 0.9 | 0.03 |
| Lower limb greater (/4) | 0.6 ± 0.8 | 0.5 ± 0.7 | 1.0 ± 1.0 | <0.01 |
| Upper limb sum (/8) | 1.9 ± 1.4 | 1.8 ± 1.4 | 2.1 ± 1.6 | 0.19 |
| Upper limb lesser (/4) | 0.7 ± 0.7 | 0.6 ± 0.7 | 0.8 ± 0.8 | 0.25 |
| Upper limb greater (/4) | 1.2 ± 0.8 | 1.2 ± 0.8 | 1.3 ± 0.9 | 0.22 |
| Neck (/4) | 1.0 ± 0.8 | 0.9 ± 0.8 | 1.2 ± 0.9 | 0.09 |

P values reflect univariate tests of central tendency (*t*-tests or chi-squared tests) between Fallers and Non-Fallers.

**Table S2.** Odds ratios (OR) describing associations between individual limb rigidity subscores and fall history in the study sample.

| Rigidity Subscore | OR | 95% CI | P value | Change from main model |
| --- | --- | --- | --- | --- |
| Lower limb lesser | 1.24 | 0.38–4.03 | 0.73 | -23.0% |
| Lower limb greater | 1.99 | 0.75–5.28 | 0.17 | +23.6% |
| Upper limb lesser | 0.70 | 0.27–1.81 | 0.46 | -1.4% |
| Upper limb greater | 0.74 | 0.31–1.72 | 0.48 | +4.2% |

P values reflect logistic regressions controlling for age, sex, total UPDRS-III, PD duration, and presence of FOG.

### Associations between rigidity and less frequent falls

In addition to the associations with frequent falls discussed in the main text, additional analyses considered associations between rigidity subscores and less frequent falls. For these analyses, participants were classified as “Fallers” if they scored ≥1 on item 12 of the gait and falls questionnaire (GF-Q),^23^ “How often do you fall?” corresponding to annual or more frequent falls, and as Non-Fallers otherwise. Multivariate logistic regressions were performed to identify associations between rigidity subscores and faller status while controlling for age, sex, total UPDRS-III, PD duration, and presence of FOG as discussed in the main text.

Among the study sample, 93/216 (43%) fell annually or more often. Multivariate logistic regression results of the model with history of annual or more frequent falls as the outcome (rather than history of monthly or more frequent falls) indicated that the association between lower limb rigidity and fall history described in the main text only holds for frequent falls. Multivariate logistic regression results (Table S3) demonstrated that the association between lower limb rigidity and history of annual or more frequent falls was significantly attenuated (- 21.7%) from that of the main model, and was no longer statistically significant (P=0.119). Changes in associations between other rigidity subscores and fall history were also substantial (11-26%).

Consistent with the literature,^3, 25^ these logistic regression models also identified significant effects of female sex (Odds Ratio [95% CI] 2.1 [1.1–4.1], P=0.031), FOG (2.5 [1.2–5.1]; P=0.016), disease duration (1.1 [1.0–1.2]; P=0.021), and total UPDRS score (1.1 [1.1–1.2]; P=0.021).

**Table S3.** Odds ratios (OR) describing associations between rigidity subscores and history of annual or more frequent falls in the study sample.

| Rigidity Subscore | OR | 95% CI | P value | Change from main model |
| --- | --- | --- | --- | --- |
| Total rigidity | 0.97 | 0.85–1.10 | 0.62 | -11.0% |
| Lower limb sum | 1.26 | 0.94–1.69 | 0.12 | -21.7% |
| Upper limb sum | 0.82 | 0.60–1.12 | 0.21 | +15.5% |
| Neck | 0.74 | 0.47–1.17 | 0.74 | -26.0% |

P values reflect logistic regressions controlling for age, sex, total UPDRS-III, PD duration, and presence of FOG.

### Associations between rigidity and fall history while controlling for the presence of postural instability

To test whether associations between rigidity and fall history described in primary models were affected by the presence of postural instability (UPDRS-III item 30, the “retropulsion test”), additional multivariate logistic regression models were calculated that controlled for this variable as well as for age, gender, PD duration, total UPDRS-III score, and presence of FOG.^25^ UPDRS-III item 30 was dichotomized as ≥2 or <2, as used in a prospective study of frequent falls in PD^5^ and entered as a dichotomous predictor variable. Among the study sample, 14/216 (6%) exhibited postural instability as defined above.

Multivariate logistic regression results of the further adjusted model indicated that the associations between rigidity subscores and fall history described in the main text hold when controlling for the presence of postural instability. Numerical results (Table S4) were essentially unchanged from those of the main model, with average changes in identified OR of 1.2%, including a slight strengthening (+3.7%) for the association between lower limb rigidity and fall history.

**Table S4.** Odds ratios (OR) describing associations between rigidity subscores and history of monthly or more frequent falls in the study sample, further adjusted for presence of Postural Instability (UPDRS-III item 30 ≥2).

| Rigidity Subscore | OR | 95% CI | P value | Change from main model |
| --- | --- | --- | --- | --- |
| Total rigidity | 1.10 | 0.94–1.29 | 0.24 | +0.9% |
| Lower limb sum | 1.67 | 1.13–2.47 | <0.01 | +3.7% |
| Upper limb sum | 0.70 | 0.45–1.09 | 0.12 | -1.0% |
| Neck | 1.01 | 0.55–1.89 | 0.96 | +1.0% |

P values reflect logistic regressions controlling for age, sex, total UPDRS-III, PD duration, presence of FOG, and presence of Postural Instability.

### Associations between rigidity and fall history while controlling for the presence of impaired gait

To test whether associations between rigidity and fall history described in primary models were affected by the presence of impaired gait (UPDRS-III item 29), additional multivariate logistic regression models were calculated that controlled for this variable as well as for age, gender, PD duration, total UPDRS-III score, and presence of FOG.^25^ UPDRS-III item 29 was dichotomized as ≥2 or <2 and entered as a dichotomous predictor variable. Among the study sample, 32/216 (15%) exhibited impaired gait as defined above.

Multivariate logistic regression results of the further adjusted model indicated that the associations between rigidity subscores and fall history described in the main text hold when controlling for the presence of gait impairment. Numerical results (Table S5) were essentially unchanged from those of the main model, with average changes in identified OR of 1.0%, including a slight reduction (-8.1%) for the association between lower limb rigidity and fall history. We note that the association between lower limb rigidity and fall history was reduced to borderline statistical significance (P=0.060) when this control was added.

**Table S5.** Odds ratios (OR) describing associations between rigidity subscores and history of monthly or more frequent falls in the study sample, further adjusted for presence of impaired gait (UPDRS-III item 29 ≥2).

| Rigidity Subscore | OR | 95% CI | P value | Change from main model |
| --- | --- | --- | --- | --- |
| Total rigidity | 1.09 | 0.92–1.28 | 0.31 | +0.0% |
| Lower limb sum | 1.48 | 0.98–2.23 | 0.06 | -8.1% |
| Upper limb sum | 0.81 | 0.51–1.84 | 0.12 | +14.1% |
| Neck | 0.97 | 0.55–1.89 | 0.96 | -3.0% |

P values reflect logistic regressions controlling for age, sex, total UPDRS-III, PD duration, presence of FOG, and presence of impaired gait.

### Associations between rigidity and fall history while controlling for overall cognition

To test whether associations between rigidity and fall history described in primary models were affected by the presence of reduced overall cognition, additional multivariate logistic regression models were calculated that controlled for MoCA score as well as for age, gender, PD duration, total UPDRS-III score, and presence of FOG^25^ using the subset of cases (N=194) for whom MoCA scores were available. MoCA score was dichotomized around the sample median as ≥25 or <25 and entered as a dichotomous predictor variable. Among cases for whom MoCA scores were available (N=194), 86 (44%) exhibited reduced overall cognition as defined above.

Multivariate logistic regression results of the further adjusted model indicated that the associations between rigidity subscores and fall history described in the main text hold when controlling for the presence of reduced overall cognition. Numerical results for total, lower limb, and neck rigidity (Table S6) were essentially unchanged from those of the main model, with average changes in identified OR of 0.7%, including a slight increase (+4.3%) for the association between lower limb rigidity and fall history. The inclusion of MoCA as a covariate substantially impacted the association between neck rigidity and fall history (-20.6% change), which may indicate some interaction between neck rigidity and cognition as risk factors for falls.

**Table S6.** Odds ratios (OR) describing associations between rigidity subscores and history of monthly or more frequent falls in the study sample, further adjusted for presence of impaired gait (UPDRS-III item 29 ≥2).

| Rigidity Subscore | OR | 95% CI | P value | Change from main model |
| --- | --- | --- | --- | --- |
| Total rigidity | 1.07 | 0.91–1.26 | 0.43 | -2.0% |
| Lower limb sum | 1.68 | 1.13–2.52 | 0.01 | +4.3% |
| Upper limb sum | 0.71 | 0.43–1.17 | 0.18 | -0.3% |
| Neck | 0.79 | 0.38–1.68 | 0.55 | -20.6% |

P values reflect logistic regressions controlling for age, sex, total UPDRS-III, PD duration, presence of FOG, and dichotomized MoCA score.

### Effect of medication state during exam on associations between rigidity and fall history

Additional analyses considered whether medication state at testing (ON vs. OFF medications) affected associations between rigidity subscores and fall history. Among the 212/216 patients for whom medication information was available, 131 were taking carbidopa/levodopa (37 as monotherapy); 88, 50, 31, and 31 were taking dopamine agonists, MAO-B inhibitors, COMT inhibitors, and amantadine, respectively. Although the majority of participants (197/216) were ON medications during UPDRS assessments, some who were prescribed antiparkinsonian medications (19/216) were assessed in the practically-defined 12-hour OFF state according to the protocol of the study in which they participated.

In order to assess whether medication state affected the main study results, additional multivariate logistic regression analyses were performed considering only those patients in the ON medication state during testing. Changes in identified OR between rigidity subscores and fall history from those calculated in the main model were compared qualitatively. Because of the small number of patients in the OFF state, multivariate logistic regressions were inappropriate among these participants.

Multivariate logistic regression results considering only those patients assessed in the ON medication state indicated that the associations between rigidity subscores and fall history described in the main text hold when limited to the ON state typically observed clinically. In particular, the association between leg rigidity and fall history was only slightly attenuated (- 4.3%) and remained statistically significant (P=0.045) at the reduced sample size (Table S7). We noted that the association between neck rigidity and fall history was increased (+22%), although confidence intervals were wide (P=0.57). This suggests a trend towards an association between ON medications neck rigidity and falls.

**Table S7.**
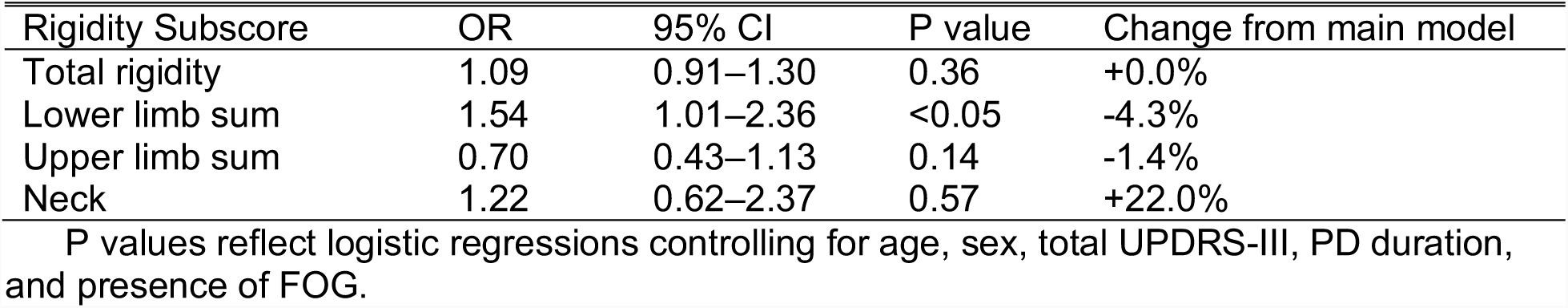
Odds ratios (OR) describing associations between rigidity subscores and history of monthly or more frequent falls among those in the study sample assessed in the ON medication state.

## References

1. Bloem BR, Grimbergen YAM, Cramer M, Willemsen M, Zwinderman AH. Prospective assessment of falls in Parkinson’s disease. J Neurol 2001;248(11):950–958.

2. Deane KH, Flaherty H, Daley DJ, et al. Priority setting partnership to identify the top 10 research priorities for the management of Parkinson’s disease. BMJ Open 2014;4(12):e006434.

3. Paul SS, Canning CG, Sherrington C, Lord SR, Close JC, Fung VS. Three simple clinical tests to accurately predict falls in people with Parkinson’s disease. Mov Disord 2013;28(5):655–662.

4. Fahn S, Elton RL, Members of the UPDRS Development Committee. The Unified Parkinson’s Disease Rating Scale. In: Fahn S, Marsden CD, Calne DB, Goldstein M, eds. Recent Developments in Parkinson’s Disease. Florham Park, NJ: Macmillan Healthcare Information, 1987:153–163.

5. Paul SS, Allen NE, Sherrington C, et al. Risk factors for frequent falls in people with Parkinson’s disease. J Parkinsons Dis 2014;4(4):699–703.

6. Latt MD, Lord SR, Morris JG, Fung VS. Clinical and physiological assessments for elucidating falls risk in Parkinson’s disease. Mov Disord 2009;24(9):1280–1289.

7. Gray P, Hildebrand K. Fall risk factors in Parkinson’s disease. J Neurosci Nurs 2000;32(4):222–228.

8. Wood BH, Bilclough JA, Bowron A, Walker RW. Incidence and prediction of falls in Parkinson’s disease: a prospective multidisciplinary study. J Neurol Neurosurg Psychiatry 2002;72(6):721–725.

9. Wright WG, Gurfinkel VS, Nutt J, Horak FB, Cordo PJ. Axial hypertonicity in Parkinson’s disease: direct measurements of trunk and hip torque. Exp Neurol 2007;208(1):38–46.

10. Horak FB, Frank J, Nutt J. Effects of dopamine on postural control in parkinsonian subjects: scaling, set, and tone. Journal of Neurophysiology 1996;75(6):2380–2396.

11. Schenkman M, Morey M, Kuchibhatla M. Spinal flexibility and balance control among community-dwelling adults with and without Parkinson’s disease. J Gerontol A Biol Sci Med Sci 2000;55(8):M441–445.

12. Franzen E, Paquette C, Gurfinkel VS, Cordo PJ, Nutt JG, Horak FB. Reduced performance in balance, walking and turning tasks is associated with increased neck tone in Parkinson’s disease. Exp Neurol 2009;219(2):430–438.

13. Weinrich M, Koch K, Garcia F, Angel RW. Axial versus distal motor impairment in Parkinson’s disease. Neurology 1988;38(4):540–545.

14. Bartolic A, Pirtosek Z, Rozman J, Ribaric S. Postural stability of Parkinson’s disease patients is improved by decreasing rigidity. Eur J Neurol 2005;12(2):156–159.

15. Zach H, Dirkx M, Pasman JW, Bloem BR, Helmich RC. The patient’s perspective: The effect of levodopa on Parkinson symptoms. Parkinsonism Relat Disord 2017;35:48-54.

16. Bingham JT, Choi JT, Ting LH. Stability in a frontal plane model of balance requires coupled changes to postural configuration and neural feedback control. Journal of Neurophysiology 2011;106(1):437–448.

17. Marusiak J, Jaskolska A, Budrewicz S, Koszewicz M, Jaskolski A. Increased muscle belly and tendon stiffness in patients with Parkinson’s disease, as measured by myotonometry. Mov Disord 2011;26(11):2119–2122.

18. Kim S, Horak FB, Carlson-Kuhta P, Park S. Postural Feedback Scaling Deficits in Parkinson’s Disease. Journal of Neurophysiology 2009;102(5):2910–2920.

19. Cano-de-la-Cuerda R, Vela-Desojo L, Miangolarra-Page JC, Macias-Macias Y. Axial rigidity is related to the risk of falls in patients with Parkinson’s disease. NeuroRehabilitation 2017;40(4):569–577.

20. van der Marck MA, Klok MP, Okun MS, et al. Consensus-based clinical practice recommendations for the examination and management of falls in patients with Parkinson’s disease. Parkinsonism Relat D 2014;20(4):360–369.

21. Factor SA, Scullin MK, Sollinger AB, et al. Freezing of gait subtypes have different cognitive correlates in Parkinson’s disease. Parkinsonism Relat D 2014;20(12):1359–1364.

22. Racette BA, Rundle M, Parsian A, Perlmutter JS. Evaluation of a screening questionnaire for genetic studies of Parkinson’s disease. Am J Med Genet 1999;88(5):539–543.

23. Giladi N, Shabtai H, Simon ES, Biran S, Tal J, Korczyn AD. Construction of freezing of gait questionnaire for patients with Parkinsonism. Parkinsonism Relat D 2000;6(3):165–170.

24. Hoops S, Nazem S, Siderowf AD, et al. Validity of the MoCA and MMSE in the detection of MCI and dementia in Parkinson disease. Neurology 2009;73(21):1738–1745.

25. McKay JL, Lang KC, Ting LH, Hackney ME. Impaired set shifting is associated with previous falls in individuals with and without Parkinson’s disease. Gait & Posture 2018;62:220–226.

26. Steiger MJ, Thompson PD, Marsden CD. Disordered axial movement in Parkinson’s disease. J Neurol Neurosurg Psychiatry 1996;61(6):645–648.

27. Prochazka A, Bennett DJ, Stephens MJ, et al. Measurement of rigidity in Parkinson’s disease. Mov Disord 1997;12(1):24–32.

28. Hammerstad JP, Elliott K, Mak E, Schulzer M, Calne S, Calne DB. Tendon jerks in Parkinson’s disease. J Neural Transm Park Dis Dement Sect 1994;8(1-2):123-130.

29. Berardelli A, Sabra AF, Hallett M. Physiological mechanisms of rigidity in Parkinson’s disease. J Neurol Neurosurg Psychiatry 1983;46(1):45–53.

30. Tatton WG, Lee RG. Evidence for abnormal long-loop reflexes in rigid Parkinsonian patients. Brain Res 1975;100(3):671–676.

31. Marusiak J, Jaskolska A, Koszewicz M, Budrewicz S, Jaskolski A. Myometry revealed medication-induced decrease in resting skeletal muscle stiffness in Parkinson’s disease patients. Clin Biomech (Bristol, Avon) 2012;27(6):632–635.

32. Petrucci MN, Diberardino LA, Mackinnon CD, Hsiao-Wecksler ET. A Neuromechanical Model of Reduced Dorsiflexor Torque During the Anticipatory Postural Adjustments of Gait Initiation. IEEE Trans Neural Syst Rehabil Eng 2018;26(11):2210–2216.

33. Peterson DS, Lohse KR, Mancini M. Relating Anticipatory Postural Adjustments to Step Outcomes During Loss of Balance in People With Parkinson’s Disease. Neurorehabil Neural Repair 2018;32(10):887–898.

34. Caetano MJD, Lord SR, Allen NE, et al. Stepping reaction time and gait adaptability are significantly impaired in people with Parkinson’s disease: Implications for fall risk. Parkinsonism Relat Disord 2018;47:32–38.

35. Fasano A, Canning CG, Hausdorff JM, Lord S, Rochester L. Falls in Parkinson’s disease: A complex and evolving picture. Movement Disorders 2017.

36. Morris ME, Menz HB, McGinley JL, et al. A randomized controlled trial to reduce falls in people with Parkinson’s disease. Neurorehabil Neural Repair 2015;29(8):777–785.

37. Sparrow D, DeAngelis TR, Hendron K, Thomas CA, Saint-Hilaire M, Ellis T. Highly Challenging Balance Program Reduces Fall Rate in Parkinson Disease. J Neurol Phys Ther 2016;40(1):24–30.

38. Li F, Harmer P, Fitzgerald K, et al. Tai chi and postural stability in patients with Parkinson’s disease. N Engl J Med 2012;366(6):511–519.

39. Chung KA, Lobb BM, Nutt JG, Horak FB. Effects of a central cholinesterase inhibitor on reducing falls in Parkinson disease. Neurology 2010;75(14):1263–1269.

40. Henderson EJ, Lord SR, Brodie MA, et al. Rivastigmine for gait stability in patients with Parkinson’s disease (ReSPonD): a randomised, double-blind, placebo-controlled, phase 2 trial. Lancet Neurol 2016;15(3):249–258.

